# Combined quantification of intracellular (phospho-)proteins and transcriptomics from fixed single cells

**DOI:** 10.1101/356329

**Authors:** Jan. P. Gerlach, Jessie A. G. van Buggenum, Sabine E.J. Tanis, Mark Hogeweg, Branco M. H. Heuts, Mauro J. Muraro, Lisa Elze, Francesca Rivello, Agata Rakszewska, Alexander van Oudenaarden, Wilhelm T. S. Huck, Hendrik G. Stunnenberg, Klaas W. Mulder

## Abstract

Environmental stimuli often lead to heterogeneous cellular responses and transcriptional output. We developed single-cell RNA and Immunodetection (RAID) to allow combined analysis the transcriptome and intracellular (phospho-)proteins from fixed single cells. RAID successfully recapitulated differentiation-state changes at the protein and mRNA level in human keratinocytes. Furthermore, we show that differentiated keratinocytes that retain high phosphorylated FAK levels, a feature associated with stem cells, also express a selection of stem cell associated transcripts. Our data demonstrates that RAID allows investigation of heterogenenous cellular responses to environmental signals at the mRNA and phospho-proteome level.

## Background

Single-cell transcriptomics approaches have revolutionized the depth of information that can be obtained from cell populations by providing detailed insights into the states of individual cells [1–6]. This is of particular interest in cell populations that comprise poorly defined cell types or cells that pass different stages of differentiation [7, 8]. Single-cell transcriptomics however faces limitations when the interest lies with specific low expressed genes, or when information about the proteome is required. Protein quantification in combination with single-cell mRNA sequencing provides a means to classify cellular subtypes, based on specific protein features, and can provide more homogenous information as the proteome is generally less prone to fluctuations than the transcriptome. To this end, transcriptomics can be combined with fluorescent antibody staining followed by FACS analysis and index sorting [9]. Such approaches are however limited by the amount of fluorescent labels available. Mass cytometry is a different approach that allows quantification of a selection of mRNAs and epitopes [10]. The great advantage of mass cytometry is the unparalleled number of cells that can be analyzed. It is however mainly suited for targeted investigations as both mRNA and protein quantifications depend on the limited number of mass labels available. Additional targeted approaches to quantify mRNAs and proteins from single cells depend on proximity ligation based protein detection [11, 12]. Recently, antibodies tagged with DNA barcodes were applied to quantify proteins through sequencing of these antibody specific tags, in combination with single-cell mRNA sequencing. CITE-seq [5] and REAP-seq [6], the techniques that make use of this approach, represent a great leap forward as large number of antibodies can be used in a single staining experiment which allows for more detailed investigation of the proteome, while also providing single-cell transcriptomics. The valuable information these techniques deliver is unfortunately still limited to cell surface proteins, as intracellular immuno-detection requires cell permeabilization and fixation. The integration of intracellular immuno-detection is however of great interest as this opens the door to measure phosphorylation events by the use of specific antibodies. Hereby, information about processes such as signal transduction could be linked to transcriptional profiles. In order to achieve intracellular (phospho-) protein detection in combination with single-cell transcriptomics, we developed single-cell <u>R</u>NA <u>a</u>nd <u>I</u>mmuno-<u>d</u>etection (RAID). RAID employs reversible fixation to allow intracellular immunostaining with <u>A</u>ntibody <u>R</u>NA-barcode <u>C</u>onjugates (ARCs) in combination with single-cell mRNA sequencing. To substantiate the potential of RAID, we turned to human keratinocytes, the epidermal cells of the skin epithelium. Keratinocytes that reside on the basal lamina are kept in a stem cell state by the combination of signaling processes, including epidermal growth factor (EGF) signaling and contact signaling through integrins [13–15]. EGF signaling is initiated by ligand binding to the epidermal growth factor receptor (EGFR) and leads to the activation of multiple downstream pathways including MAPK and AKT signaling. Furthermore, integrins play an important role for sensing the local environment by contacting components of the extracellular matrix [14]. A central step of integrin signal transduction is the activating phosphorylation of focal adhesion kinase (FAK), which controls cellular functions including proliferation, migration and survival [16]. Keratinocyte differentiation is guided by the attenuation of integrin and EGF signaling and the upregulation of other pathways, including Notch signaling [17]. The cells gradually migrate upwards in the skin as they differentiate until they form the protective, cornified layer of the skin, which is marked by heavy crosslinking of the extracellular matrix and loss of nuclei [14]. Keratinocytes can be readily cultured as a monolayer, providing a simple system to study their differentiation *in vitro* [18].

Using the keratinocyte system we show that ARC-based intracellular protein quantification could be integrated in an adapted CELseq2 protocol for single-cell transcriptomics. The RAID staining and fixation protocol is compatible with the generation of mRNA libraries of high gene complexity, despite a mild diminution of the gene detection rate. Finally, we performed RAID to assess the response of keratinocytes to EGFR inhibition. According to our expectations, both ARC and mRNA measurements illustrated that keratinocytes transitioned from a stem cell to differentiated state after EGFR inhibition. EGFR inhibition leads to a reduction of FAK phosphorylation at the population level. Interestingly, ARC-based quantifications also revealed that this response was highly heterogeneous and some cells retained high levels of phosphorylated FAK after differentiation. We show that these FAK-retaining cells express elevated levels of several stem cell marks, including integrin substrates. EGFR inhibition therefore heterogeneously affects the integrin pathway at the mRNA and signaling level. We conclude that RAID successfully combines transcriptomics with ARC based measurements of intracellular and extracellular epitopes at single-cell resolution.

## Results

We designed RAID to combine quantification of intracellular (phospho-) proteins and the transcriptome from individual cells in a streamlined workflow (Figure 1, a more detailed overview is depicted in Figure S1). Two key challenges needed to be addressed for the development of RAID. First, protein measurements needed to be incorporated in the library preparation during single-cell RNA profiling. We solved this issue through the use of Antibody RNA-barcode Conjugates (ARCs) in combination with a modified CELseq2 procedure [19, 20]. Second, intracellular immunostaining requires permeabilization of the cells to allow antibodies to cross the plasma membrane. Cell permeabilization without cell fixation would however lead to loss of the endogenous mRNAs. Application of a fixation strategy that is compatible with single-cell RNA-sequencing therefore was required. This issue was resolved by performing the cell fixation with the combination of two chemically reversible crosslinking reagents that allow the release of both the endogenous RNA as well as the antibody-RNA conjugates. The final RAID procedure essentially consists of three main steps. 1) Reversible cell fixation and immunostaining with antibody-RNA conjugates; 2) reverse crosslinking and first strand cDNA synthesis; and 3) single-cell cDNA samples are pooled and a sequencing library is generated that comprises both mRNA and ARC libraries (Figure 1). Detailed RAID immunostaining and sequencing-library preparation protocols are provided as Additional Files FI and F2, respectively.

**Figure 1.**
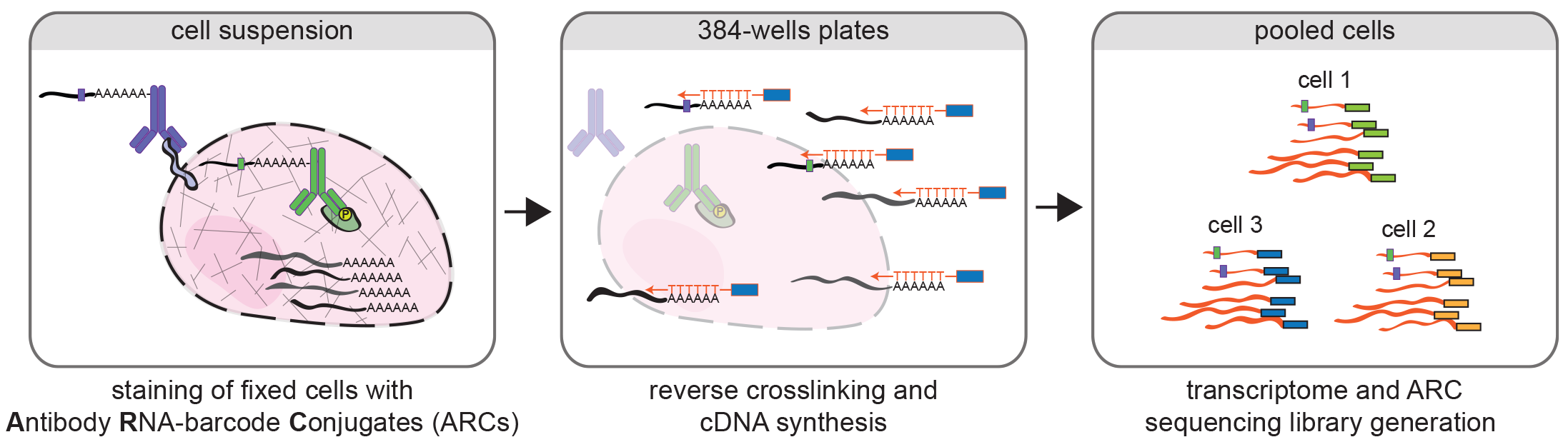
Schematic overview of the RAID workflow. Cells are crosslinked, permeabilized and stained with Antibody RNA-Barcode Conjugates (ARCs) in suspension. Hereafter, cells are sorted into 384-wells plates where crosslinking is reversed and cDNA synthesis is performed using CELseq2 compatible dTV primers. Finally, single-cell samples are pooled and sequencing library preparation is performed using an adapted CELseq2 protocol to incorporate ARC measurements in the library.

## Simultaneous quantification of antibody-RNA conjugates (ARCs) and the transcriptome by RNA-sequencing

To implement RAID, we first needed to establish robust detection of both the ARCs as well as the endogenous transcriptome from single cells by using a modified SORT-seq/CELseq2 procedure [19, 20]. The RNA-barcodes we constructed contain a 10 base-pair antibody-specific barcode and a unique molecular identifier (UMI). This allows multiplexed count-based quantification of antibody signals, principally similar to previous applications [5, 6, 21]. Moreover, these RNA-barcodes are 5’ capped and 3′ poly-adenylated to resemble endogenous mRNAs that can be incorporated into a single-cell RNA-sequencing library preparation workflow. For a detailed overview of the RNA-barcode design see Figure S2 and the materials and methods section.

Experiments using bulk human keratinocyte cultures showed that epidermal growth factor receptor (EGFR) inhibition results in downregulation of EGFR protein levels and a slight upregulation of integrin-α6 (ITGA6) protein levels (Figure S3A). As inhibition of EGFR induces epidermal differentiation [18, 21], the ratio of these cell surface proteins consequently provides information about the differentiation state of the cells (Figure 2A). Because cell surface staining does not require fixation and permeabilisation, we first focused on the detection of ARCs for the EGFR and ITGA6 from unfixed keratinocytes as a proof of principle. We prepared sequencing libraries of ARC stained keratinocytes from untreated cells and from cells that were treated with the EGFR inhibitor AG1478 for 48 hours to induce differentiation [17, 18]. After sequencing, read mapping and UMI-based molecular counting, ARCs and endogenous transcripts were readily detected from the same cells (Figure 2B and Figure S3B).

**Figure 2.**
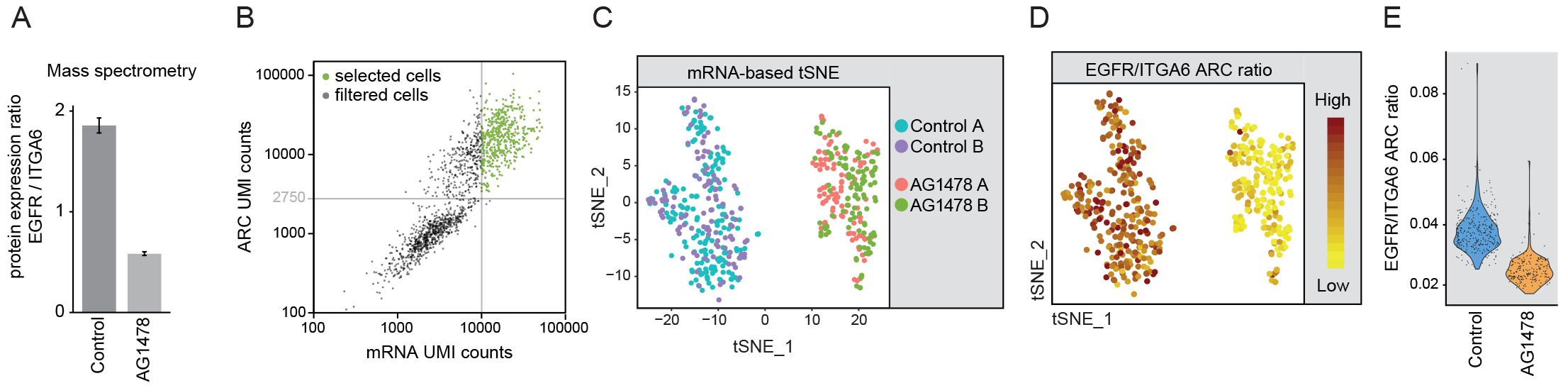
Combined single-cell transcriptomics and antibody-barcode conjugate (ARC) detection from unfixed keratinocytes. (A) Boxplot showing mass spectrometry based analysis of the EGFR to ITGA6 ratio from bulk samples. AG1478 induced differentiation leads to a reduction of the EGFR to ITGA6 ratio. Experiment was performed in triplicate. (B-C) Single-cell analysis of unfixed keratinocytes stained with ARCs for EGFR and ITGA6. Cell were untreated or treated with the EGFR inhibitor AG1478 to induce differentiation. (B) Scatterplot shows the total number of UMI counts that were detected per cell from the transcriptome and from ARCs. Cells that passed the count thresholds for the mRNA (10000) and ARCs (2750) are indicated in green. (C) tSNE visualization using principal component 1 to 8 of the variable transcripts. Untreated and AG1478 treated cells form distinct clusters. Different batches of cells are color coded. (D) Featureplot showing the ratio of the UMI counts of EGFR-ARC and ITGA6-ARC for each cell projected on the tSNE coordinates from C. (E) Ratio of the UMI counts of EGFR-ARC and ITGA6-ARC for control and AG1478 treated cells represented in violin plot.

To investigate whether our data recapitulates expected differentiation changes at both the mRNA and protein level, we selected high quality cells with more than 2,750 ARC counts and 10,000 mRNA counts. Selected cells covered a median of 4594 detected genes per cell (range: 2,811 - 8642 genes, Figure 2B and S3B). After normalizing for differences in sequencing depth by subsampling, we performed principal component analysis (PCA) and subsequent t-distributed stochastic neighbor embedding (tSNE) visualization on the single-cell transcriptomes. This revealed separate clusters for the untreated and AG1478 treated cells, while the different batches/plates from the same treatment intermingled (Figure 2C). Assessment of selected stem cell and differentiation markers [22] at the transcriptome level revealed that these clusters indeed represent differentiated (AG1478 treated) and undifferentiated (untreated) cell states, respectively (Figure S3C and S3D, [22]). Moreover, differential expression and gene ontology (GO) overrepresentation analysis confirmed that AG1478 treatment indeed resulted in upregulation of genes involved in epidermal cornification and keratinocyte differentiation (Figure S3E). Next, the ratio of EGFR over ITGA6 levels was calculated for each individual cell, based on the ARC measurements, showing that the EGFR to ITGA6 ratio is indeed reduced in AG1478 treated cells compared to non-treated samples (Figure 2D and 2E). Consistent with the protein measurements, EGFR mRNA expression is downregulated in response to AG1478, both at the single-cell, as well as the population level (Figures S3F, S3G, S3H). In contrast, ITGA6 mRNA remains largely unaffected by AG1478 (Figure S3I, S3J, S3K) suggesting that its protein levels are regulated post-transcriptionally. Taken together, these experiments establish that our antibody-RNA conjugates and modified CELseq2 sample preparation approach enable combined protein and mRNA quantification from single cells.

## High quality RNA-sequencing of reversibly fixed and permeabilized single-cells

RAID was designed to allow quantification of intracellular (phosphorylated) epitopes, which requires fixation and permeabilization. Generating high quality single-cell RNA-sequencing profiles from such samples however is challenging as the recovery of endogenous RNA could be affected by crosslinking or by diffusion from the cells after permeabilization. To achieve sufficient fixation to prevent loss of RNA during immunostaining, yet allow efficient retrieval of RNA for library preparation, RAID uses a combination of the chemically reversible crosslinker DSP (dithiobis(succinimidyl propionate)), which has previously been shown to be compatible with mRNA sequencing [23], and the related crosslinker SPDP (succinimidyl 3-(2-pyridyldithio)propionate). DSP crosslinks proteins by its two NHS groups that are reactive towards primary amines as found in lysines, while SPDP contains a single NHS group and a maleimide moiety that reacts with sulfhydryls, most commonly found in cysteines that are not engaged in disulfide bonds. The two reactive groups of DSP and SPDP are separated by a disulfide-containing linker, which can be cleaved by reducing agents (e.g. DTT). For efficient simultaneous release of the RNA-barcodes during reverse crosslinking, we also introduced a disulfide-bond in the linker connecting the RNA-barcodes to the antibodies.

To assess to what extend the RAID workflow influences the quality of single-cell mRNA sequencing, we performed a direct comparison of unfixed cells with cells that underwent the RAID procedure, including mock antibody staining. After sequencing, we detected a median of 6,180 genes per cell in the unfixed samples based on 40,000 subsampled reads (n=62 cells, Figure 3A). Even though RAID cells displayed a reduced number of genes detected (5,552 genes per cell, n=53 cells, Figure 3A), the achieved gene complexity is expected to allow detailed analysis of the transcriptome. Furthermore, the reduced gene complexity in RAID cells appears to be caused by a subtle overall reduction of the gene detection rate (Figure 3B) indicating that fixation, cell permeabilization and antibody staining does not result in loss of specific sets of RNAs. In agreement, the average gene expression profile was highly similar between unfixed and RAID cells (R^2^=0.90, Figure 3C). Our experiments show that the RAID procedure, including reversible fixation, permeabilization and mock antibody staining generates high quality single-cell mRNA sequencing data.

**Figure 3.**
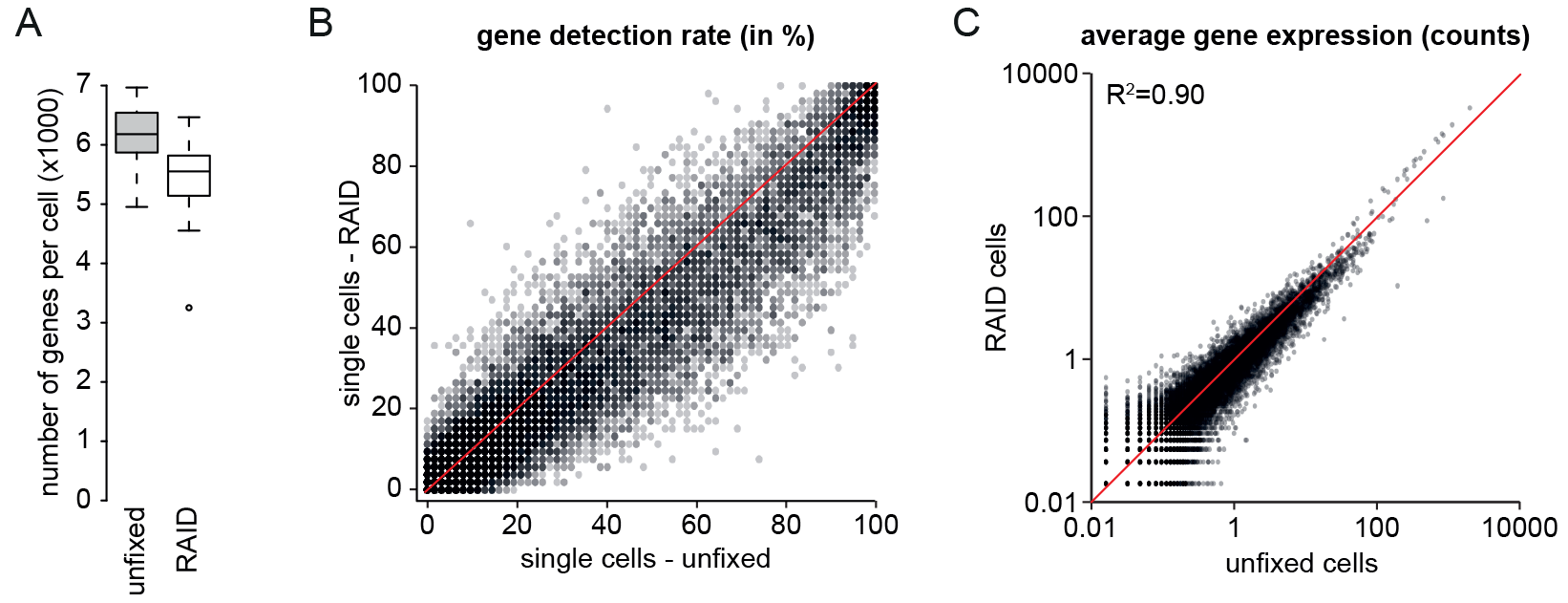
Effect of the RAID procedure on single-cell transcriptomics quality. Comparison of single-cell transcriptomics quality between unfixed cells and cells that passed the complete RAID procedure including DSP/SPDP based fixation and mock staining. Comparison is based on 40000 sampled UMI counts per cell. (A) Boxplots showing a mild reduction of the gene complexity after the complete RAID procedure. (B) Scatterplot showing the detection rate of genes in unfixed cells and RAID cells. (C) Scatterplot showing the average expression of genes in unfixed cells and RAID cells. (B,C) Red line represents the x=y function as a visual aid.

## Phosphorylated FAK is associated with the retention of stem cell marks during keratinocyte differetiation

Human epidermal keratinocyte differentiation involves loss of EGFR and integrin mediated signaling and activation of others, including the Notch pathway [17]. Moreover, progression of differentiation is associated with changes in transcriptional programs to accommodate new functions of the cells. We generated ARCs to monitor EGFR (phosphorylated RPS6) and integrin (phosphorylated FAK) pathway activity during differentiation. RPS6 is phosphorylated downstream of active EGF signaling [21] and is lost within the first 24 hours of AG1478 treatment (Figure S4A). Quantification of phospho-RPS6-ARCs therefore serves as an indicator of EGFR signaling in individual keratinocytes. Integrin signaling cooperates with EGFR signaling to maintain keratinocytes in their stem cell state [13, 14]. Focal adhesion kinase (FAK) is a key component of integrin signaling and its activating phosphorylation (pFAK) plays a central role in keratinocyte biology. Furthermore, we included two ARCs directed against components of the Notch signaling pathway (NOTCH1 and JAG1), which plays a role in the onset of differentiation [17]. Finally, ARCs targeting kallikrein-6 (KLK6) and transglutaminase-1 (TGM1), factors involved in extracellular matrix remodeling and cornification during differentiation, were included as exemplars of canonical differentiation markers [24, 25]. The selected antibodies were previously verified and used in an antibody-DNA barcode staining platform developed in our lab (ID-seq) [21].

We used this panel of ARCs in a RAID experiment with untreated cells and cells that were treated for 48 hours with AG1478 to induce differentiation. High-quality transcriptome and ARC profiles were obtained from n=630 control cells and n=515 AG1478 treated cells. As anticipated, PCA and subsequent tSNE visualization of the mRNA data showed a clear separation of control and AG1478 cells (Figure 4A). Moreover, expression of specific markers [22], as well as GO enrichment analysis, confirmed AG1478-induced differentiation (Figure S4B-D). RAID-ARC measurements revealed a reduction of pRPS6 and pFAK signals upon AG1478 treatment, indicating effective inhibition of EGFR activity, decreased integrin-mediated signaling and onset of differentiation (Figure 4B and 4C). This was confirmed by the observation that the differentiation associated proteins NOTCH1, TGM1 and KLK6 were significantly upregulated in these cells (Figure 4B and C, Kolmogorov-Smirnov test, p<0.01). In contrast, JAG1 levels were slightly, but significantly reduced in the AG1478 sample (Figure 4B and C, Kolmogorov-Smirnov test, p<0.01). The data indicates that the RAID-ARC measurements recapitulate the expected differentiation-induced dynamics observed in bulk samples (Figure S4E).

**Figure 4.**
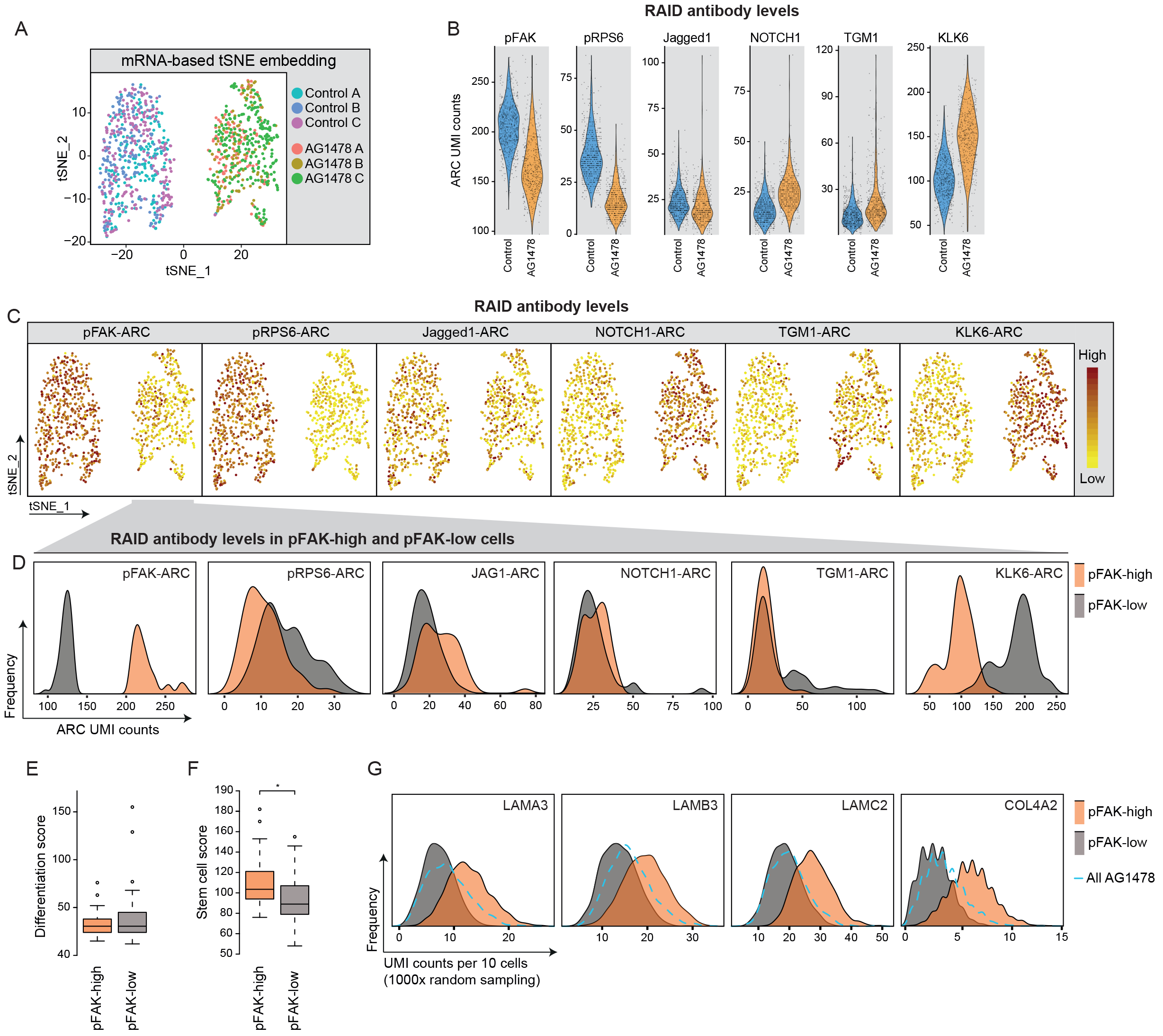
RAID analysis shows association of FAK phosphorylation with the expression of stem cell marks. RAID was performed with ARC-based quantification of six antibodies from fixed keratinocytes. Cells were untreated or treated with the EGFR inhibitor AG1478 to induce differentiation. (A) tSNE visualization using principal component 1 to 8 of the variable transcripts. Untreated and AG1478 treated cells form distinct clusters. Different batches of cells are color coded. (B) Violin plots representing the ARC UMI counts for the indicated antibodies, based on a total of 400 sampled ARC counts per cell. (C) Featureplots showing data from B, projected on the tSNE embeddings from A. (D-G) From the AG1478 treated cells, 50 cells were selected with the highest and lowest pFAK-ARC counts (pFAK-high and pFAK-low, respectively) and analyzed based on proteomic and transcriptomic features. (D) Histograms showing the distribution of UMI counts for the indicated ARCs in pFAK-high (orange) and pFAK-low cells (grey). (E) Boxplots showing the differentiation scores of pFAK-high and pFAK-low cells. (F) Boxplots showing the stem cell scores of pFAK-high and pFAK-low cells. The stem cell score is significantly higher in pFAK-high cells based on two tailed t-test (p=0.0023). (G) Histograms showing distribution of UMI counts of the indicated integrin substrates per ten cells after lOOOx random sampling from pFAK-high (orange), pFAK-low cells (grey) and total AG1478 treated cells (blue dashed line). Transcript UMI counts are based on 4500 sampled UMI counts per single cell.

A challenge for single-cell techniques is to not only capture the global differences between distinct groups of cells, but to also reveal biological heterogeneity within cell populations. The spread in ARC levels suggests that we captured considerable heterogeneity in the cellular response to AG1478 treatment. This is exemplified by the observation that a part of AG1478-treated cells preserve phospho-FAK (pFAK) levels comparable to those in untreated cells (Figure 4B and 4C). Due to the central role of FAK in integrin signaling in keratinocytes, we sought to investigate this population in more detail. To this end, we selected 50 cells with the highest- and lowest- phospho-FAK levels from the AG1478-treated cell population (pFAK-high and pFAK-low, respectively), and analyzed the differences between these cells based on the other measured proteins, as well as the transcriptome. Interestingly, pFAK-low cells show higher KLK6-ARC signals and a fraction of these cells also exhibits elevated TGM1-ARC counts compared to differentiated cells that retain FAK phosphorylation (Figure 4D, Kolmogorov-Smirnov test, p<0.05). Furthermore, JAG1 levels are somewhat reduced in pFAK-low cells (Figure 4D, Kolmogorov-Smirnov test, p<0.05). Interestingly, pFAK-high cells displayed lower pRPS6 signals than pFAK-low cells, indicating that differences between these populations did not arise from inefficient EGFR inhibition by AG1478 (Figure 4D). Significant changes in NOTCH1-ARC levels were not detected. Based on these combined observations we asked if pFAK-low and pFAK-retaining cells could encompass different states of differentiation.

To investigate this, we turned to the matched transcriptome data and asked if pFAK-retaining cells harbor stem cell characteristics in their transcriptome. To obtain an unbiased measure of sternness and differentiation, we calculated a stem cell and a differentiation score for each cell by aggregating the UMI counts of 530 stem cell associated mRNAs and 226 differentiation-associated mRNAs, respectively ([22] and Additional File F3). Comparing untreated and AG1478 treated samples indicates that the stem cell score is elevated in the undifferentiated cells, whereas the differentiation score is strongly increased in AG1478 treated cells (Figure S5A-C). This demonstrates that these mRNA-based scores reflect the cellular differentiation state of the epidermal keratinocytes.

We found that the mean differentiation scores of pFAK-high and pFAK-low cells are equal, indicating that both populations are in a similar differentiated state based on their RNA-expression programs (Figure 4E). In contrast, the stem cell score is significantly higher in pFAK-high cells compared to pFAK-low cells (Figure 4F, 2-tailed t-test, p<0.01). Thus, pFAK-high cells retain stem cell associated transcripts at higher levels, while already expressing differentiation genes at comparable levels to fully differentiated cells. To gain some insight into why these populations may display these differences we performed differential expression analysis, revealing that 15 of the 531 stem cell associated transcripts were expressed at significantly higher levels in pFAK-high cells compared to pFAK-low cells (Additional File F4, 2-tailed t-test, p<0.05). Strikingly, these 15 stem cell markers included four excreted substrates of integrin receptors (LAMA3, LAMB3, LAMC2, COL4A2). To visualize the mRNA levels of these integrin substrates in pFAK-high cells, we randomly sampled 10 cells from either pFAK-high or pFAK-low cells 1000 times and analyzed the sum of the UMI counts from each of the differentially expressed laminins and collagens. Each of these genes showed increased expression in pFAK-high cells compared to pFAK-low cells, as well as to cells randomly sampled from all AG1478 treated cells (Figure 4G). We conclude that cells that fail to downregulate phosphorylated FAK levels, despite AG1478-induced differentiation, retain additional stem cell features including transcripts associated with integrin signaling. Taken together, our experiments show that RAID does not only allow detection of global changes between cell populations, based on their differentiation state, but can also provide information about heterogeneous responses at the transcriptional and (phospho-) proteomic level.

## Discussion

The development of techniques that combine single-cell transcriptomics with protein measurements is a great step for the characterization of cells from heterogeneous populations [5, 6]. Protein quantifications can take into account post-transcriptional and post-translational regulation and thereby provide more comprehensive information about cellular regulation when integrated with transcriptomic information. This is exemplified in our system by the upregulation of ITGA6 after AG1478 induced differentiation despite a subtle reduction of mRNA levels (Figures S3B and S3H). Furthermore, biologically relevant transcripts are often expressed at the limit of detection. In some cases, antibody based protein quantification can therefore provide information about markers that are expressed at too low levels for robust detection at the mRNA level. For example, in our experiments we were unable to quantify the differentiation markers TGM1 and KLK6 at the mRNA level while ARC measurements clearly showed the expected upregulation of the proteins after differentiation. We anticipate that, especially when shallow sequencing of large numbers of cells is performed, the addition of marker proteins can greatly aid the grouping and characterization of the cells. The applicability of this approach was previously demonstrated with CITE-seq and REAP-seq using cell surface markers to categorize blood cells [5, 6]. Although the protein quantification of cell surface markers provides valuable information, especially in the blood system, inclusion of intracellular epitopes that could reveal information on the cell state in terms of intracellular proteins and their phosphorylation, has not been described. The RAID technology now enables these analyses.

We developed the RAID library generation using a plate-based system and a modified CELseq2/SORT-seq protocol [19, 20], whereas the related techniques REAP-seq and CITE-seq make use of droplet microfluidics and a separate library prep to generate libraries for antibody tags and mRNA [5, 6]. The key advantage of microfluidic platforms is the high number of cells that can be processed with relatively little effort. However, plate-based systems in combination with sorting have the distinct advantage of facilitating pre-selection to discard unwanted materials such as cell doublets or debris, while specific cells could be enriched based on specific markers. Furthermore, plate-based systems are capable of producing single-cell libraries of high gene complexity, as we also demonstrate here, which allows very detailed investigation of the transcriptome. As reverse crosslinking of DSP/SPDP is based on chemical reduction, which was shown to be compatible with microfluidic processing [6], we anticipate that the RAID is readily transferrable to such systems.

Using a panel of six antibodies, we showed that RAID enables the quantification of extracellular and intracellular proteins, including two phosphorylated epitopes, each representing relevant processes during keratinocyte differentiation. The changes that we detected with each of the ARCs during differentiation resembled the changes of independent experiments in bulk samples, indicating that they provide biologically accurate information. Besides these global changes at the protein level that were detected between stem cells and differentiated cells, it is clear that each population contains a certain amount of heterogeneity in the ARC signals. By zooming in on cells that retained high levels of phosphorylated FAK after differentiation we showed that several integrin substrates were upregulated in pFAK-high cells. Based on our data, it is unclear whether the expression of the integrin substrates represents a cause or consequence of high pFAK levels. Nonetheless, differentiation initiated by EGFR inhibition heterogeneously affects the integrin pathway at the mRNA and signaling level. Another factor that is potentially linked to the heterogeneous response of pFAK is the differentiation marker KLK6. KLK6 is a protease that targets multiple proteins within the extracellular matrix. Interestingly, the laminin family of integrin substrates are among its targets [26] and pFAK-high cells showed a striking reduction of KLK6. KLK6 levels could therefore negatively modulate Integrin signaling and pFAK levels. Further investigation is however required to substantiate these speculations. Taken together, our results illustrate that RAID can be applied to link transcriptional and proteomic changes to specific phosphorylation events.

## Conclusion

Single cell RAID provides a means to perform single cell transcriptomics in combination with antibody-RNA conjugate-based measurements from fixed single cell. The RAID workflow only leads to a minor reduction of gene complexity and preserves biological differences between cell populations. The RAID fixation procedure allows the detection of intracellular epitopes, including phosphorylation events on proteins. Finally, by focusing on differentiated cells that retain high levels of phosphorylated FAK we demonstrate that RAID allows analysis of heterogeneous responses to extracellular cues at the level of the phospho-proteome and transcriptome. We anticipate that RAID can provide important insights in the connection between signaling pathways and the transcriptome in the future.

## Declarations

### Ethics approval and consent to participate

“Not applicable”

### Consent for publication

“Not applicable”

### Availability of data and material

The raw and processed datasets generated during the current study are available under the GEO accession number GSE115981.
[https://www.ncbi.nlm.nih.gov/geo/query/acc.cgi?acc=GSE115981]

## Authors’ contributions

JG and KM conceived the method. Single cell experiments were performed by JG and BH. JvB developed the RAID analysis pipeline. JG, JvB and BH analysed and interpreted data. Antibody-RNA conjugations were set-up and performed by JG, JvB, MH and LE. ST performed the proteomics experiments and mRNA-seq experiments from bulk samples. Setup of the adapted CELseq/SORT-seq procedure MM, AO and JG. FR, AR and WH contributed reagents and expert advice. HS and KW oversaw the study. JG, HS and KWM wrote the manuscript with input from all authors.

## Acknowledgements

The authors thank Peter Brazda and Eva Janssen-Megens for help with the preparation of sorting plates and sequencing and Rob Woestenenk for help with FACS sorting.

## Materials and Methods

### Cell culture and AG 1478 treatment

Human keratinocytes (pooled foreskin strain, Lonza) were cultured as described [21]. After expansion of the keratinocytes on J2-3T3 feeder cells, the keratinocytes were grown for 2 days on keratinocyte serum-free medium (KSFM) with supplements (30 μg/mL bovine pituitary extract and 0.2 ng/mL EGF) (Gibco). For induction of differentiation, KSFM was supplemented with 10 μM AG1478 (Calbiochem) 48 hours before cell collection.

### RNA-barcode production

RNA barcodes were produced by *in vitro* transcription with the mMessage mMachine T7 IVT kit from Ambion using 100-500 ng template DNA in 10 μl reactions with the addition of 0.5 μl of RNAsin Plus (Promega). The DNA template design is (5’-<u>GGATCCTAATACGACTCACTATAGGG</u>AGACCGACGAAACTGTTAACGTCGCACGACGC-TCTTCCGATCT<u>GTCAGTCA</u>**NNNNNNNNNNNNNNN[<u>l0 nt antibody barcode</u>]**ATCAGTCAACAGATAAGCGTGAGATAG-GGCATTACCGAGGCCTGGAGCATTGCCGATACCGAGAGTATTAGCTACGTTGCAGAGGATGCGACGGATGCAAAAAAAAAAAAAAAAAAAAAAAAAAAA). The DNA templates contain a T7 promoter sequence (underlined), a stagger sequence with the of length of 1-8 nucleotides (italic, underlined), a 15 nucleotide UMI, and an antibody-specific barcode sequence. The specific barcode sequences used for our experiments are indicated in Additional Figure 1. After IVT, the RNA-barcodes were purified using the Microprep RNA purification kit from Zymo Research and eluted in 10 μl water. Typically yields of purified RNA barcodes are between 10 to 20 μg.

### RNA-barcode functionalization and conjugation to antibodies

The strategy of RNA to antibody conjugation is based on similar chemical principles as previously described for antibody DNA conjugation [27]. To this end, antibodies and nucleotide barcodes are functionalized with the respective reactive groups tetrazine and trans-cyclooctene that allow highly efficient conjugation through the inverse electron-demand Diels-Alder reaction [28]. First, purified RNA barcodes were 3’ end-labelled with pCp-Azide using T4 RNA ligase in an overnight reaction of 10 μl at 16°C. The reaction mixtures were composed of (3 ng RNA-barcode, 1X T4 RNA ligase buffer (New England Biolabs), 1 mM ATP (New England Biolabs), 10 % DMSO, 0.1 mM pCp-Azide (Jena Bioscience), 0.5 μl T4 RNA Ligase (New England Biolabs) and 0.5 μl RNAsin Plus (Promega)). After the reaction, excess pCp-Azide was removed by RNA purification with the Microprep RNA purification kit from Zymo Research. Next, purified RNA-Azide was 3’ end-labelled with trans-Cyclooctene by incubating overnight with a 20-fold molar excess of DBCO-PEG12-TCO (trans-Cyclooctene-PEG12-Dibenzylcyclooctyne, Jena Bioscience) at room temperature (e.g. 3 μg RNA-Azide in 15 μl ultrapure water with 62.5 μM DBCO-PEG12-TCO). Hereafter, the RNA-TCO (trans-Cyclooctene labelled RNA) was purified with the Microprep RNA purification kit from Zymo Research to dispose of excess DBCO-PEG12-TCO. RNA-TCO was eluted with water and taken up in 50 mM borate buffered saline pH 8.4. Antibodies in 50 mM Borate Buffered Saline pH 8.4 were functionalized with NHS-SS-Tetrazine as described [21, 27]. 1.5 μg (= 21.5 pmol) functionalized RNA-Barcodes (RNA-TCO) were conjugated to 5 μg (= 33.5 pmol) functionalized antibodies by incubation in Borate Buffered Saline pH 8.4 at room temperature for 1 hour in the presence of 0.5 U/μl of RNAsin Plus.

### Antibodies

All antibodies were ordered without any carrier proteins that could interfere with antibody functionalization. EGFR (Ab231, Abcam), NOTCH1 (AF5317, RnD systems), JAG1 (AF1277, RnD systems), KLK6 (AF2008, RnD systems), phospho-FAK (AF4528, RnD systems), phospho-RPS6 (5364, Cell Signaling Technology), ITGA6 (Produced and purified in house from hybridoma P5G10, DSHB), TGM1 (Produced and purified in house from BC1 hybridoma, a kind gift from Prof. Robert Rice), alpha-tubulin (T6074, Sigma-Aldrich).

### Single-cell RAID - fixation and immunostaining

A detailed protocol for the fixation and Immunostaining procedure is provided in Additional File F1. Keratinocytes were collected by trypsinization and taken up in Sodium Phosphate Buffered Saline pH 8.4. Cells were fixed using a combination of 2.5 mM DSP (Thermo Scientific) and 2.5 mM SPDP (Thermo Scientific) for 45 minutes in Sodium Phosphate Buffered Saline pH 8.4. After fixative quenching with [100 mM Tris-HCl pH 7.5, 150 mM NaCl] the cells were blocked and permeabilized using [0.5 × Protein Free Blocking Buffer (Thermo scientific) in PBS, supplement with 100 μg/ml of pre-boiled tRNA (Roche), 0.5 U/μl RNAsin Plus (Promega) and 0.1 % Triton X100]. For ARC staining of unfixed cells as presented in Figure 2, the fixation step was skipped and Triton X100 was excluded from the blocking buffer. Next, cells were stained overnight with ARCs in a staining buffer containing [0.5 × PFBB in PBS, supplement with 2 U/μl RNAsin Plus (Promega), 0.1 % Triton X100 and 250 ng/μl of each Antibody RNA-Barcode Conjugate]. Triton X100 was excluded from the buffer for staining of unfixed cells and staining was reduced to two hours. After immunostaining, the cells were gently washed 6 times with 10 ml wash buffer [0.1X PFBB in PBS] and transferred to FACS tubes.

### Single-cell RAID - library preparation

A detailed protocol for the RAID library preparation is provided in Additional File F2. The RAID library prep is based on the SORT-seq/CELseq2 library prep [19, 20], with the indicated adaptations. Single cells were sorted in 384-wells plates containing 100 nl of (7.5 ng/μl) unique CELseq2 compatible primers in the wells and 5 μl mineral oil (Sigma-Aldrich). These reverse transcription primer sequences were adapted to allow sequencing of the transcripts/ARC sequences in read 1 and the cell barcode and UMI in read 2 (Additional File F5). After sorting, the plates were frozen at −80 °C and thawed before further processing. Reverse crosslinking was performed by dispensing 50 nl of a 3X reverse-crosslinking buffer [6 mM dNTP, 150 mM Tris pH 8, 90 mM DTT, 0.1 % Triton X100, 6 U/μl RNAsin Plus (Promega)] into the wells and incubating 45 minutes at 25 °C. Hereafter, the reactions were incubated 5 minutes at 65 °C and cooled to 4 °C. Next, reverse transcription was performed in a total reaction volume of 250 nl using the Maxima H minus Reverse transcriptase (Thermo Scientific). To enhance second strand synthesis of the ARC sequences, 50 nl of 0.3 pmol/μl Barcode Compensation Primer [5′ GGGAGACCGACGAAACTGTTAACG] solution was dispensed into the wells. The samples were heated to 85 °C for 5 minutes and the slowly cooled down by decreasing the temperature 5 degrees every 30 seconds until reaching 10 °C. Next, the samples were pooled and purified. Second strand synthesis and *in vitro* transcription were performed as described [21]. Reverse transcription of the amplified RNA was performed with Maxima H minus Reverse Transcriptase (Thermo Scientific) using a combination of a random octamer primer [5’ C ACG ACG CT CTT CCGATCTNNNNNNNN] and the Barcode Compensation Primer [5’ GGGAGACCGACGAAACTGTTAACG] for enhanced priming of ARC sequences. Library preparation PCR was performed in two steps. First a library pre-amplification with short primers [Forward 5’ C ACG ACG CT CTT CCG AT CT, Reverse 5’ GTTCAGACGTGTGCTCTTCCGATC] was performed using the Herculase II enzyme (Agilent) to minimize amplification bias. Next, adapter extension was performed using a PCR reaction with Herculase II (Agilent) and the following primers (Forward Library primer: [5’ AATGATACGGCGACCACCGAGATCTACACTCTTTCC-CTACACGACGCTCTTCCGATCT], Reverse indexing Primer [5′ CAAGCAGAAGACGGCATACGAGAT**[6nt-index]**GTGACTGGAGTTCAGACGTGTGCTCTTCCGATC]). Sequencing was performed using the NextSeq500 from Illumina with 63 bases for read 1, 14 bases for read 2 and 6 bases for the Illumina index. Note that for the comparison unfixed and RAID cells, as presented in Figure 3, the cells were sorted into 96 wells plates containing two sets of 48 unique CELseq2 compatible primers and the reaction mixtures for the reverse transcription were scaled to 2 μl. Raw and processed datasets from our experiments (count tables) have been deposited under GEO accession number GSE115981.

### Data analysis and representation

Sequence data was demultiplexed using bcl2fastq software (Illumina). Transcriptome count tables were generated with the published CELseq2 pipeline [20], using the following settings [min_bc_quality = 10, cutjength = 50,]. To allow compatibility with the pipeline, the read names for read 1 and read 2 were swapped. ARC count-tables were generated using a combination of the CELseq2 pipeline and adapted version of the ID-seq pipeline which was previously published for quantification of the DNA-version of our antibody-barcode conjugates [21]. First, all reads were assigned to specific cells using the demultiplexing function of the CELseq2 pipeline in conjunction with the transcriptome analysis. Next, the ARC sequences were retrieved from the reads and counted using an adapted version of ID-seq pipeline [21], which is available at https://github.com/jessievb/RAID. In this approach, ARC sequences are identified using a 12 nucleotide identification sequence within the barcode [5’ ATCAGTCAACAG], allowing one mismatch. Next, the 10 nt antibody specific sequences and the 15 nt UMIs are extracted and listed. Note that in this approach the total nucleotide length for specific antibody identification spans 22 nucleotides. Finally, UMIs for each antibody specific barcode were counted and a count table is generated listing all cells and antibodies. In our approach, ARC and transcriptome count tables contain identical cell names which allows for facile combination of the two datasets.

For read-depth normalization of the transcriptomes and selection of high quality cells, a fixed number of UMI counts was randomly sampled from each cell and cells with fewer counts than this threshold were discarded from the analysis. We used a count threshold of 10000 UMI count for the cell surface stained cells (Figure 2), 40000 for the comparison of unfixed and RAID cells (Figure 3) and 4500 for the RAID experiment with intracellular staining (Figures 3 and 4). ARC-stained cells were also filtered for a minimum number of ARC counts (2750 counts for cell surface stained cells, Figure 2 and 400 counts for RAID cells, Figure 4). ARC counts for the RAID experiment were normalized by sampling 400 ARC counts per cell.

Further processing of the mRNA and ARC data was performed using the Seurat R package [29]. First, the transcriptome data (subsampled count tables) was Log normalized. Next, variable genes were defined and used for PCA. tSNE was performed using principal components 1 to 8. Subsampled ARC count tables, or ARC ratios were loaded into the Seurat object as dataset for multi-modal analysis. As subsampling the ARC data or calculation of the ARC ratios corrects for differences in overall ARC abundances between cells, no additional normalization was performed. To reduce outlier effects in featureplot representations, the 5% and 95% percentiles of the data were set to the minimum and maximum of the color scale, respectively. Differential expression analysis was performed using the FindMarkers function of the Seurat package using a log fold-change threshold of 0.25. Gene Ontology analyzation of differentially expressed genes was performed using goseq [30].

### CELseq2 for bulk samples

Sequencing library generation was performed according to the CELseq2 protocol with minor adaptations, as described [20, 21]. Purified RNA was added into 96 wells plates containing CELseq2 compatible primers. Reverse transcription was performed in 2 μl reactions overlaid with 7 μl Vapor-Lock (Qiagen) using the Maxima H minus reverse transcriptase (ThermoFisher). Further steps were performed as described [21]. Sequencing was performed using the NextSeq500 from Illumina.

### Western blotting

Keratinocytes were grown in culture plates and directly lysed using 1X SDS sample buffer containing 1 % SDS, 50mM TrisHCI pH 6.8, 10 % glycerol, 50 mM DTT and Bromophenol blue. Samples were boiled and centrifuged for 1 minute at 16,000 xg. Samples were separated on mini-PROTEAN 4-20% TGX gels (Biorad) and blotted on PVDF membranes. Membranes were blocked with Protein Free Blocking Buffer (Thermo scientific) and Immunostained overnight with phospho-RPS6 (5364, Cell Signaling Technology) and alpha-tubulin (T6074, Sigma-Aldrich) antibodies. Staining with secondary antibodies (LiCor) was performed for one hour and membranes where scanned using the Odyssey CLx imaging system from LiCor.

### Mass-spectrometry of bulk samples

Cells were harvested, washed, snap frozen and stored at −80 °C. Cells were lysed by boiling in a lysis buffer containing 4 % SDS, 100 mM Tris-HCl pH 7.6, 100 mM DTT for 3 minutes at 95 °C. DNA was sheared using sonication. The samples were centrifuged for 5 minutes at 16.000 × g at 4 °C and the supernatant was taken for protein quantification with the PierceTM BCA Protein Assay Kit (Thermo Scientific). For the generation of tryptic peptides, we applied filter-aided sample preparation [31]. For absolute quantification the proteins we used a standard range of proteins (UPS2-1SET, Sigma) which we spiked into one of the samples (3,3 μg in sample equivalent to 100.000 cells). To obtain deep-proteome, samples were fractionated using strong anion exchange, collecting fractions of the flow through and elutions at pH 11, 8, 5 and 2 of Britton and Robinson buffer. Samples were desalted and concentrated using C18 stage-tips [32]. The peptide samples were separated on an Easy nLC 1000 (Thermo Scientific) connected online to a Thermo Scientific Orbitrap Fusion Tribrid mass spectrometer. A 240 min acetonitrile gradient (5-23 %, 8-27 %, 9-30 %, 11-32 % and 14-32 % for FT, pH 11, 8, 5 and 2, respectively) was applied to the five fractions. MS and MS/MS spectra were recorded in a Top speed modus with a run cycle of 3s. MS/MS spectra were recorded in the Ion trap using Higher-energy Collision Dissociation fragmentation. To analyze the raw mass spectrometry data we used MaxQuant (version 1.5.1.0, database: Uniprot_201512\HUMAN) [33] with default setting and the “match between runs” and “iBAQ” algorithms enabled. We filtered out reverse hits and imputed missing values using Perseus (default settings, MaxQuant software package).

**Figure SI. Detailed overview of the RAID procedure.** Cells are collected and crosslinked in suspension using the reversible crosslinkers DSP and SPDP. Hereafter, cells are permeabilized using 0.1X Triton X100 and blocked using Protein Free Blocking buffer. Immunostaining with Antibody RNA-Barcode Conjugates (ARCs) is performed overnight. After extensive washing of the cells, they are sorted into 384-wells plates containing unique CELseq2 compatible primers, overlaid with mineral oil. To allow efficient lysis and reverse crosslinking, cells are first frozen at −80, followed by the addition of a DTT containing reverse crosslinking buffer. Next, reverse transcription is performed which incorporates a specific barcode in the cDNA from each cell and therefore allows sample pooling. Sequencing library preparation is performed according to an adapted CELseq2 protocol to allow efficient incorporation of ARC signals. The final sequencing library is composed of a broad mRNA signal and a specific ARC peak.

**Figure S2. Sequence overview of the RAID antibody RNA-barcodes.** The RNA barcode contains a 5’ Cap which is incorporated during the *in vitro* transcription based barcode production with the goal to enhance barcode stability and resemblance to mRNAs. The second strand enhancer is a sequence that is used to enhance priming of the second strand synthesis for RNA-barcodes. The Illumnia adapter is part of the complete adapter sequence required for sequencing. The stagger sequence is a sequence of barcode dependent length that aims to prevent sequence failure due to overrepresentation of a common barcode sequence. The UMI within the RNA-barcode is use for the quantification of the ARCs. The antibody barcode sequence is the antibody specific sequence that allows antibody identification and ARC multiplexing. The RNA-barcode includes an extension sequence that enhances the efficiency of the CELseq2 based library prep for the ARCs. Finally, the RNA-barcodes contain a 28nt polA tail to mimic cellular mRNAs and allow CELseq2 based library preparation.

**Figure S3. Additional data to Figure 2, Combined single-cell transcriptomics and Antibody RNA-barcode Conjugate (ARC) detection from unfixed keratinocytes.** (A) Boxplots showing the estimated copy numbers of EGFR and ITGA6 proteins in mass-spectrometry based proteomics of untreated and AG1478 treated Keratinocytes. The experiment was performed in triplicate. (B) Scatterplot showing the number of genes detected per cell, related to the total UMI counts of the transcriptome. Cells that passed the count thresholds for the mRNA (10000) and ARCs (2750) are indicated in green. (C) Featureplots showing the normalized mRNA expression of a selection of stem cell markers projected on the tSNE coordinates from Figure 2C. (D) Featureplots showing the normalized mRNA expression of a selection of differentiation markers projected on the cells of the tSNE coordinates from Figure 2C. (E) GO analysis shows biological processes significantly upregulated after AG1478 induced differentiation. (F, G) Normalized EGFR mRNA expression projected on the tSNE embedding from Figure 2C (F) and represented in violin plot (G). (H) Boxplots showing the EGFR mRNA expression in bulk analysis of untreated and AG1478 treated keratinocytes. The experiment was performed in triplicate. (I, J) Normalized ITGA6 mRNA expression projected on the tSNE embeddings from Figure 2C (I) and represented in violin plot (J). (K) Boxplots showing the ITGA6 mRNA expression in bulk analysis of untreated and AG1478 treated keratinocytes. The experiment was performed in triplicate.

**Figure S4. Additional data to Figure 4. RAID analysis shows association of FAK phosphorylation with the expression of stem cell marks.** (A) AG1478 treatment triggers loss of phosphorylated RPS6 as shown by western blotting and immunodetection. (B) GO analysis shows biological processes significantly upregulated in RAID cells after AG1478 induced differentiation. (C) Featureplots showing the normalized mRNA expression of a selection of stem cell markers projected on the cells of the tSNE from Figure 4A. (D) Featureplots showing the normalized mRNA expression of a selection of differentiation markers projected on the cells of the tSNE from Figure 4A. (E) Boxplots showing the estimated cellular copy numbers of TGM1, KLK6, JAG1 and NOTCH1 proteins in mass-spectrometry based proteomics of untreated and AG1478 treated keratinocytes. The experiment was performed in triplicate. Significant differences between untreated and AG1478 treated cells are indicated by asterisk (2-tailed t-test p<0.05).

**Figure S5. Stem cell and differentiation scores summarize the differentiation status of keratinocytes.** (A) Stem cell and differentiation scores projected on the tSNE plot from Figure 4A. (B) Boxplots showing the stem cell scores of untreated and AG1478 treated RAID cells (C) Boxplots showing the differentiation scores of untreated and AG1478 treated RAID cells.

